# Tirzepatide attenuates atherosclerosis through weight loss-independent anti-inflammatory mechanisms

**DOI:** 10.64898/2026.06.22.733886

**Authors:** Shuo Chen, Shulin Wei, Tian Tian, Zhenghong Liu, Meiming Su, Fanshun Zhang, Yanjun Yin, Meijie Chen, Lin Jiang, Paul C Evans, Bradford C. Berk, Stefan Offermanns, Yihai Cao, Zhihua Wang, Jianping Weng, Suowen Xu

## Abstract

**Background:** Atherosclerosis is a chronic inflammatory vascular disorder with persistent residual inflammation even after standard lipid-lowering therapy. Mounting evidence from bench to bedside suggests that diabetes and obesity accelerate atherosclerosis development. Tirzepatide (TZP), a dual Glucagon-Like Peptide-1 Receptor/Glucose-Dependent Insulinotropic Polypeptide Receptor (GLP-1R/GIPR) agonist approved for treating diabetes and obesity, has demonstrated proven cardiometabolic efficacy in large cardiovascular outcome trials. However, it remains largely uncertain whether TZP attenuates atherosclerosis independent of its anti-diabetic and anti-obese effects through direct actions on the vasculature.

**Methods:** We established atherosclerotic mouse models under diabetic, obese, and non-diabetic/non-obese conditions. Analysis of covariance (ANCOVA) and pair-feeding experiments were applied to experimentally decouple weight-dependent metabolic improvement from intrinsic vasculoprotection. Molecular and cell biological assays in human umbilical vein endothelial cells (HUVECs) and human aortic endothelial cells (HAECs) were performed to dissect the underlying signaling mechanisms.

**Results:** TZP markedly reduced aortic plaque burden and inflammation, restrained necrotic core enlargement, and improved plaque stability across all experimental mouse models. Both ANCOVA and pair-feeding experiments confirmed that these atheroprotective effects were independent of food intake and body weight loss. Furthermore, TZP attenuated systemic and vascular inflammation in Tumor Necrosis Factor-α (TNF)-treated C57BL/6J mice, and this protection occurred without changes in body weight or blood glucose levels. Mechanistically, TZP directly targeted endothelial cells and activated the cyclic adenosine monophosphate (cAMP)/protein kinase A (PKA)/endothelial nitric oxide synthase (eNOS) pathway, increased eNOS phosphorylation and nitric oxide bioavailability, consequently downregulating the expression of the pro-inflammatory adhesion molecules Vascular Cell Adhesion Molecule-1 (VCAM–1) and Intercellular Adhesion Molecule-1 (ICAM–1).

**Conclusions:** TZP arrests atherosclerosis progression through weight loss-independent anti-inflammatory mechanisms. These findings implicate TZP as a promising therapeutic drug for mitigating residual vascular inflammation in patients with atherosclerotic cardiovascular disease (ASCVD), irrespective of glycemic status or obesity.

**Clinical Perspective:** *What Is New?:*

- Tirzepatide exerts direct anti-atherosclerotic effects in preclinical mouse models of atherosclerosis under diabetic, obese, and non-obese conditions.
- Tirzepatide directly targets endothelial GLP-1R/GIPR and downstream cAMP/PKA/eNOS signaling pathway to suppress NF-κB-driven vascular inflammation, thereby uncovering a previously unrecognized vasculoprotective mechanism underlying its cardiovascular benefits

*What Are the Clinical Implications?:*

- Tirzepatide exerts direct vascular protective effects independent of body weight reduction, suggesting that its cardiovascular benefits may extend beyond glycemic control and obesity management.
- Tirzepatide may represent a promising therapeutic drug for addressing residual vascular inflammation in ASCVD patients, including those without overt diabetes or obesity

## Introduction

Excess caloric intake and a sedentary lifestyle have led to a global epidemic of obesity and diabetes. These metabolic disorders, in turn, contribute substantially to the rising morbidity and mortality of atherosclerotic cardiovascular disease (ASCVD), imposing an increasingly heavy burden on global public health systems^1^. Metabolic dysfunction promotes the initiation and progression of atherosclerosis, the shared pathological basis of ASCVD, by driving interconnected pathological processes, including chronic inflammation, dysregulated adipokine signaling, and insulin resistance^2–4^.

Since dysregulated lipid metabolism is currently recognized as a primary driver of atherosclerosis progression, lipid-lowering therapy represents the cornerstone of its clinical prevention and treatment. However, lowering blood lipids alone does not entirely eliminate cardiovascular risk, as growing epidemiological and clinical evidence underscores the critical role of vascular inflammation in driving plaque progression^5–7^. Crucially, chronic low-grade inflammation often persists even after effective control of metabolic risk factors such as dyslipidemia and hyperglycemia, thereby continuing to drive atherosclerosis progression^8,9^. Thus, targeting residual inflammation, in addition to metabolic management, may provide a promising therapeutic strategy.

Metabolic disorders such as diabetes and obesity accelerate inflammatory response in panvascular tissues. In this respect, Tirzepatide (TZP), an U.S. Food and Drug Administration (FDA)-approved dual Glucagon-Like Peptide-1/Glucose-Dependent Insulinotropic Polypeptide (GLP-1/GIP) receptor agonist, lowers blood glucose and body weight as well as systemic inflammation associated biomarkers, and reduces major cardiovascular events in patients with diabetes and obesity^10–13^. However, it remains unresolved whether TZP’s anti-inflammatory and cardiovascular actions are secondary to weight loss and metabolic improvement–or reflect a direct vasculoprotective effect^14–17^. This question is clinically important: if TZP directly suppresses vascular inflammation independent of weight loss, it could benefit a broader population, including lean individuals with residual inflammation despite optimal lipid and metabolic control, and guide more targeted use of GLP-1/GIP-based therapies.

Here, we investigated whether TZP confers direct, weight-independent protection against vascular inflammation and atherogenesis, and further delineated the underlying molecular mechanisms. Using a comprehensive panel of murine atherosclerotic models spanning diabetic, obese and non-diabetic non-obese background, we demonstrate that TZP exerts anti-atherosclerotic effects that are dissociable from its systemic weight-lowering and metabolic benefits. We further define the endothelial cell-intrinsic actions of TZP and unravel the core mechanisms mediating its vasculoprotective activity. Collectively, our findings uncover a previously unappreciated, weight loss-independent vascular protective dimension of TZP, establishing a novel mechanistic framework to support the repurposing of TZP to mitigate residual cardiovascular risk irrespective of diabetes or obese milieu.

## Methods

### Data Availability

All supporting data are available from the corresponding author upon reasonable request. Detailed descriptions of materials and methods are in the **Supplemental Material**. The raw sequence data of transcriptome and single-cell RNA sequencing reported in this article have been deposited in the Genomic Sequence Archive (GSA:https://ngdc.cncb.ac.cn/gsa), and their access numbers are CRA043567 and CRA043608, respectively.

### Experimental mice and treatment

Eight-week-old male *Apoe^-/-^* mice and C57BL/6J were purchased from GemPharmatech (Strain NO. T001458). Eight-week-old male *ob/ob* mice were purchased from GemPharmatech (Strain NO. T001461). Hepatic Low-Density Lipoprotein Receptor (LDLR) protein expression was downregulated by a single intravenous injection of adeno-associated virus serotype 8 (AAV8)-Pcsk9D377Y (3 × 10¹¹ vg per mouse). The AAV vector was purchased from Vigene Biosciences. All mice were fed a high-cholesterol diet (HCD, Research Diets, catalog No. D12108C; containing 20% fat and 1.25% cholesterol) to induce atherosclerosis. Mice were housed in SPF-grade standard cages with a 12-hour light/dark cycle, free access to standard feed, and at an ambient temperature of 20–24 °C and humidity of 30–70%. For drug intervention, mice were randomly divided into control and TZP treatment groups. Mice in TZP treatment groups were subcutaneously administered TZP at 1 nmol/kg (low dose) or 10 nmol/kg (high dose) three times per week; control animals received an equal volume of vehicle via the same route. This intermittent subcutaneous dosage regimen (1 and 10 nmol/kg, thrice weekly) was referenced to published rodent pharmacodynamic literatures^18,19^. The animal experiment protocol has been approved by the Animal Ethics Committee of the University of Science and Technology of China (USTCACUC27120124102) and has been conducted in accordance with the guidelines for the husbandry and use of laboratory animals issued by the National Institutes of Health (NIH). All mouse experimental procedures were performed in accordance with the ARRIVE guidelines 2.0 (Animal Research: Reporting of In Vivo Experiments).

### Statistical Analysis

Data were expressed as mean ±SEM unless stated otherwise. Normality of data was verified via the Shapiro–Wilk test, and the Brown–Forsythe test was applied to evaluate homogeneity of variances prior to statistical comparison. Two-group comparisons of normally distributed data with equal variance were conducted using unpaired t-test; Welch’s t-test was adopted for normally distributed data with unequal variances, whereas nonparametric Mann–Whitney U test was used for non-normally distributed data. For comparisons across three or more experimental groups with normally distributed and homoscedastic data, one-way analysis of variance (ANOVA) was performed, with Bonferroni or Dunnett’s post hoc multiple comparison tests executed. When data satisfied normality but exhibited heterogeneous variances, Welch’s one-way ANOVA coupled with Games–Howell post hoc test was utilized instead. Experiments incorporating two independent variables were analyzed by two-way ANOVA, complemented with Tukey’s multiple comparison post hoc test. All statistical computations were completed using GraphPad Prism 10 (GraphPad Software, La Jolla, CA, USA). Two-tailed P values < 0.05 were defined as statistically significant. Representative microscopic/experimental images were chosen to represent the mean phenotypic characteristics of respective experimental cohorts.

## Results

### TZP attenuates atherosclerosis in diabetic *Apoe^-/-^* mice

We used *Apoe^-/-^* mice treated with low-dose of streptozotocin (STZ) to establish a preclinical model of diabetes-associated atherosclerosis^20^. *Apoe^-/-^* mice were fed a high-cholesterol diet (HCD) for 4 weeks and then administered with low-dose STZ to induce hyperglycemia, thereby modeling hyperglycemia-accelerated plaque formation in vivo. Following an increase and subsequent stabilization of plasma glucose levels, diabetic atherosclerosis mice were administrated low (1 nmol/kg)/high (10 nmol/kg) doses of TZP or saline, while continuing the HCD for an additional 8 weeks (**Fig. 1A**). Body weight and fasting blood glucose were well matched among all groups at baseline.

**Fig. 1.**
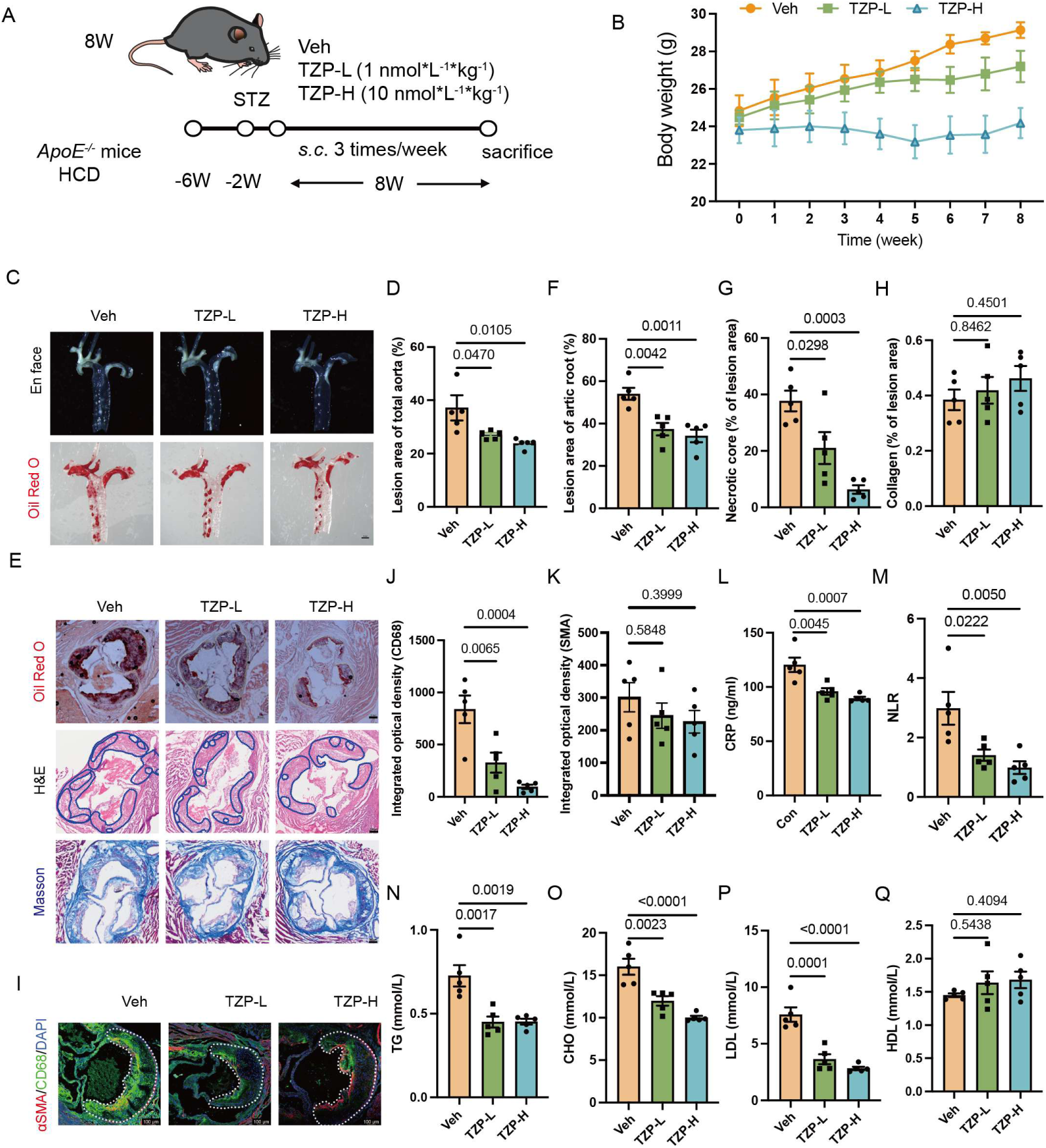
TZP attenuates atherosclerosis in diabetic *Apoe^-/-^* mice. **(A)** Schematic diagram illustrating the overall experimental design. **(B)** Longitudinal changes in body weight of diabetic *Apoe^-/-^* mice administered with vehicle (Veh), low-dose tirzepatide (TZP-L), or high-dose tirzepatide (TZP-H). **(C–D)** Representative images and quantitative analysis of en face Oil Red O staining for atherosclerotic lesions in the entire aorta. Scale bar: 1 mm. **(E–H)** Representative histological images and corresponding quantitative statistics of aortic root sections stained with Oil Red O, hematoxylin and eosin (H&E), and Masson’s trichrome staining. Scale bar: 200 μm. **(I–K)** Representative immunofluorescence images and quantitative analysis of alpha-smooth muscle actin (α-SMA, red) and cluster of differentiation 68 (CD68, green) in aortic root atherosclerotic plaques. Scale bar: 20 μm. **(L–M)** Serum C-reactive protein (CRP) levels and neutrophil-to-lymphocyte ratio (NLR). TZP treatment ameliorated systemic inflammation in diabetic *Apoe^-/-^* mice. **(N–Q)** Quantification of serum lipid profiles, including triglyceride (TG), total cholesterol (CHO), low-density lipoprotein cholesterol (LDL-C), and high-density lipoprotein cholesterol (HDL-C). Data are presented as mean ± SEM, n = 5 per group. Statistical analysis was performed using two-way ANOVA with Tukey’s post hoc test for panel B, and one-way ANOVA with Bonferroni correction for panels C–K.

Systematic glucose metabolic parameters were assessed to evaluate the effect of TZP on diabetic atherosclerosis. Both low- and high-dose TZP markedly lowered body weight in a dose-dependent manner (**Fig. 1B**) and decreased cumulative food intake (**Fig. S1A–S1C**). Consistent with this reduction in body weight, TZP-treated mice also showed decreased weight of liver, inguinal white adipose tissue (iWAT), epididymal white adipose tissue (eWAT), while showed no reduction in the weight of brown adipose tissue (BAT) (**Fig. S1E–S1G**). Importantly, TZP inhibited atherosclerosis lesion area. Oil Red O staining of the aorta revealed a significant reduction in atherosclerotic plaque area in both low-dose and high-dose treatment groups compared with controls (**Fig. 1C–1D**). Consistent with these *en face* aortic findings, Oil Red O staining of aortic sinus sections also demonstrated markedly decreased plaque burden following TZP administration. Similarly, TZP effectively reduced plaque area in the aortic sinus. H&E staining further revealed reduced necrotic core size in TZP-treated mice, whereas Masson’s trichrome staining showed no significant change in plaque collagen content (**Fig. 1E–1H**). CD68 immunofluorescence staining revealed a significant reduction in the infiltration of CD68⁺ macrophages within lesions, suggesting attenuation of plaque inflammation after TZP treatment. By comparison, α-smooth muscle actin (α-SMA) immunofluorescence quantification showed that the abundance of α-SMA-positive vascular smooth muscle cells did not differ significantly among groups (**Fig. 1I** – **1K**). Given the critical role of inflammation in atherosclerosis, we next assessed systemic inflammatory markers. Both low- and high-dose of TZP significantly decreased serum C-reactive protein (CRP) level and the neutrophil-to-lymphocyte ratio (NLR) (**Fig. 1L–1M**). Concurrently, analysis of lipid profiles showed that TZP downregulated serum triglycerides (TG), total cholesterol (TC), and low-density lipoprotein cholesterol (LDL-C), while high-density lipoprotein cholesterol (HDL-C) levels remained unchanged (**Fig. 1N–1Q**).

### TZP ameliorates atherosclerosis in obese atherogenic mice

Given the weight-lowering effects of TZP and the strong epidemiological as well as mechanistic links between obesity and plaque progression/instability^21^, we next asked whether TZP provides vascular protection in obesity-driven atherosclerosis. To address this question, we established an obesity-associated atherosclerosis model by delivering AAV-mPCSK9 to *ob/ob* mice, thereby generating a co-morbid setting mirroring obesity, insulin resistance and atherogenic dyslipidemia (**Fig. 2A**).

**Fig. 2.**
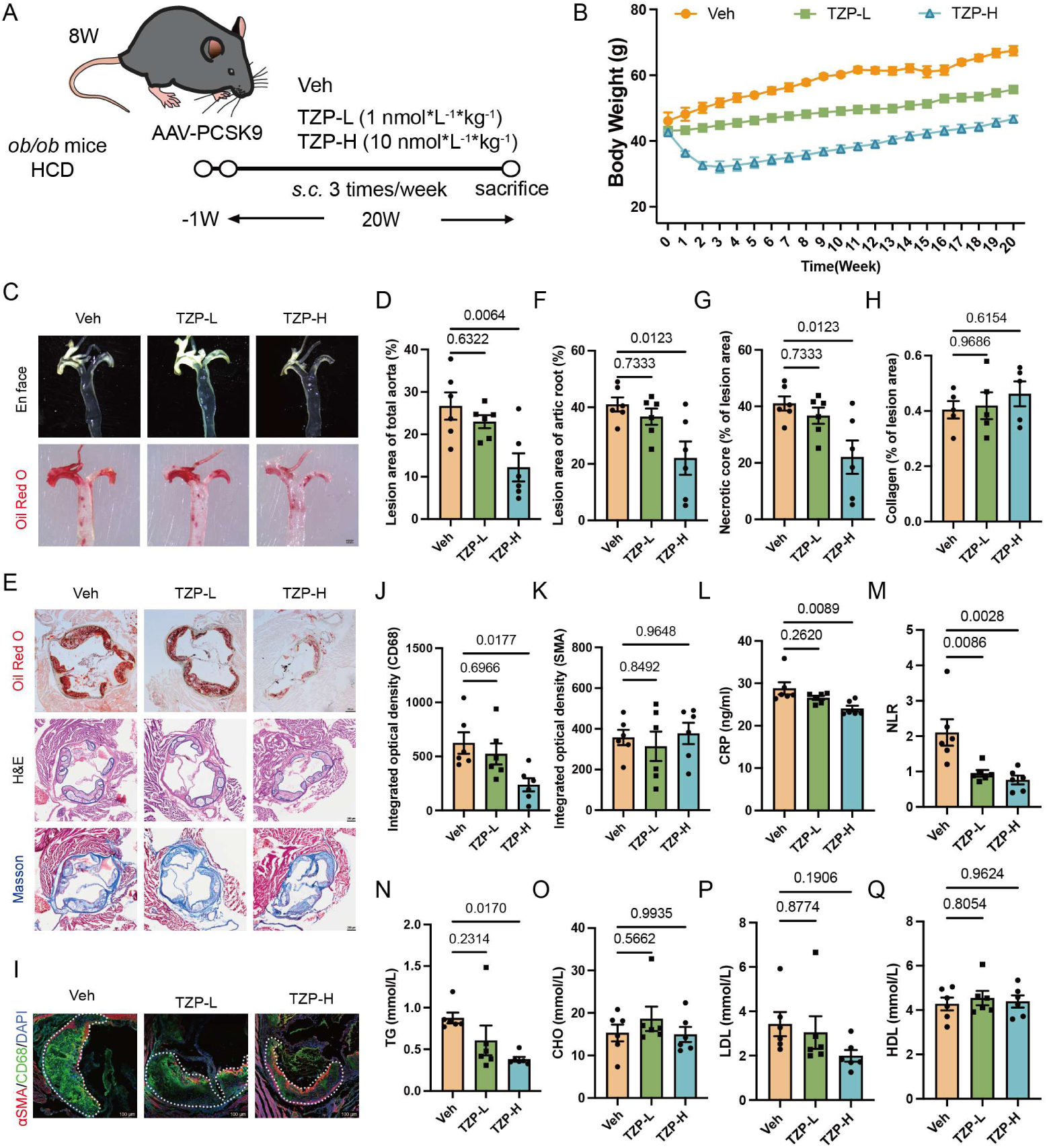
TZP attenuates atherosclerotic progression and mitigates metabolic and inflammatory disturbances in obese atherogenic mice. **(A)** Schematic overview of the animal experimental design. **(B)** Dynamic changes in body weight among obese atherogenic mice administered vehicle (Veh), low-dose tirzepatide (TZP-L), or high-dose tirzepatide (TZP-H). **(C–D)** Representative images and quantitative analysis of atherosclerotic lesions via en face Oil Red O staining of whole aortas. Scale bar: 1 mm. **(E–H)** Representative histological staining and quantitative results of aortic root sections, including Oil Red O, hematoxylin and eosin (H&E), and Masson trichrome staining. Scale bar: 200 μm. **(I–K)** Representative immunofluorescence images and corresponding quantification of aortic root plaques stained for alpha-smooth muscle actin (α-SMA; red) and cluster of differentiation 68 (CD68; green). Scale bar: 20 μm. **(L–M)** Detection of serum C-reactive protein (CRP) concentrations and neutrophil-to-lymphocyte ratio (NLR), indicating that TZP treatment alleviated systemic inflammation. **(N–Q)** Serum lipid profiles containing triglyceride (TG), total cholesterol (TC), low-density lipoprotein cholesterol (LDL-C), and high-density lipoprotein cholesterol (HDL-C) in experimental mice. Data are presented as mean±SEM; n=6 for all experimental groups. Two-way ANOVA followed by Tukey multiple comparisons test was used for statistical analysis in (B). One-way ANOVA with Bonferroni’s post hoc test was applied for (C–K).

Compared with vehicle group, TZP treatment resulted in a reduction in cumulative food intake and body weight in *ob/ob* mice (**Fig. 2B and S2A–2C**). This weight loss was accompanied by decreases in iWAT, eWAT, and the liver-to-body weight ratio, but not in BAT (**Fig. S2H–S2J**). Consistent with data in diabetic atherosclerosis mice, *en face* Oil Red O staining showed a significant reduction in aortic plaque area in the high-dose TZP-treated groups, while low-dose TZP showed no significant effect (**Fig. 2C–2D**). Further analysis on aortic sinus sections demonstrated decreased lesion area in mice receiving high-dose TZP. H&E staining identified diminished necrotic core size within plaques, while quantification of Masson’s trichrome staining indicated that plaque collagen content was not significantly altered between groups (**Fig. 2E–2H**). Immunofluorescence staining revealed reduced CD68⁺ macrophage accumulation within the plaques, while α-SMA expression increased following TZP administration (**Fig. 2I–2K**), suggesting a shift toward a less inflammatory and more stable plaque phenotype. Similarly, systemic inflammatory markers, including CRP and NLR, were markedly decreased in the high-dose TZP group, demonstrating alleviated systemic inflammation compared with control mice (**Fig. 2L–2M**).

TZP also improved glucose metabolism in the obese atherosclerosis model. Fasting glucose levels did not differ significantly between the high- or low-dose TZP treated group and control group. However, both low- and high-dose TZP treatment improved glucose tolerance and insulin sensitivity (**Fig. S2D–S2G**). At the end of the study, plasma levels of TC, HDL-C, and LDL-C were unchanged after TZP treatment, whereas TG levels were significantly reduced in the high-dose TZP treatment group (**Fig. 2N–2Q**).

In summary, in obese mice, low-dose TZP exerted limited influence on body weight and improved glucose metabolism, but had no significant impact on lipid metabolism, inflammatory status, or atherosclerosis progression. In contrast, high-dose TZP treatment effectively improved systemic inflammation and lipid metabolism, and attenuated atherosclerosis progression.

### TZP reduces atherosclerosis in non-diabetic and non-obese *Apoe^-/-^* mice

Having established the anti-atherosclerotic effects of TZP in diabetic- and obesity-associated atherosclerosis mouse models, we further examined its pharmacological effects in non-diabetic, non-obese atherosclerosis using *Apoe^-/-^* mice (**Fig. 3A**). Consistent with the data in the aforementioned two models, TZP treatment lowered body weight in *Apoe^-/-^* mice compared to control group (**Fig. 3B**) Simultaneously, assessment of the aorta revealed a marked reduction in atherosclerotic plaque burden after TZP treatment (**Fig. 3C–3D**). This protective effect was further supported by Oil Red O staining of the aortic sinus. Plaque composition in TZP-treated mice was also improved, with decreased necrotic core size and increased collagen deposition, as assessed by H&E and Masson’s trichrome staining (**Fig. 3E–3H**). Plaques from TZP-treated mice displayed a significant reduction in CD68⁺ macrophage content and an increase in α-SMA^+^ smooth muscle cell content, suggesting attenuated plaque inflammation and improved plaque stability (**Fig. 3I–3K**). To further evaluate the inflammatory status of these mice, we measured systemic inflammatory markers. Notably, *Apoe^-/-^* mice treated with TZP exhibited a significant reduction in CRP level and NLR compared with control mice, indicating reduced systemic inflammation (**Fig. 3L–3M**) Meanwhile, TZP-treated mice also showed decreased plasma TC, TG, and LDL-C levels, whereas HDL-C levels were not significantly altered (**Fig. 3N–3Q**). Similarly, TZP treatment decreased both cumulative food intake and average daily food intake (**Fig. S3A–S3C**), accompanied by body weight loss as well as decreases in weight of iWAT, eWAT, and the liver-to-body weight ratio, but BAT weight remained unchanged (**Fig. S3D-3G**). In this non-diabetic, non-obese atherosclerosis mouse model, TZP did not affect blood glucose levels (**Fig. S3C**).

**Fig. 3.**
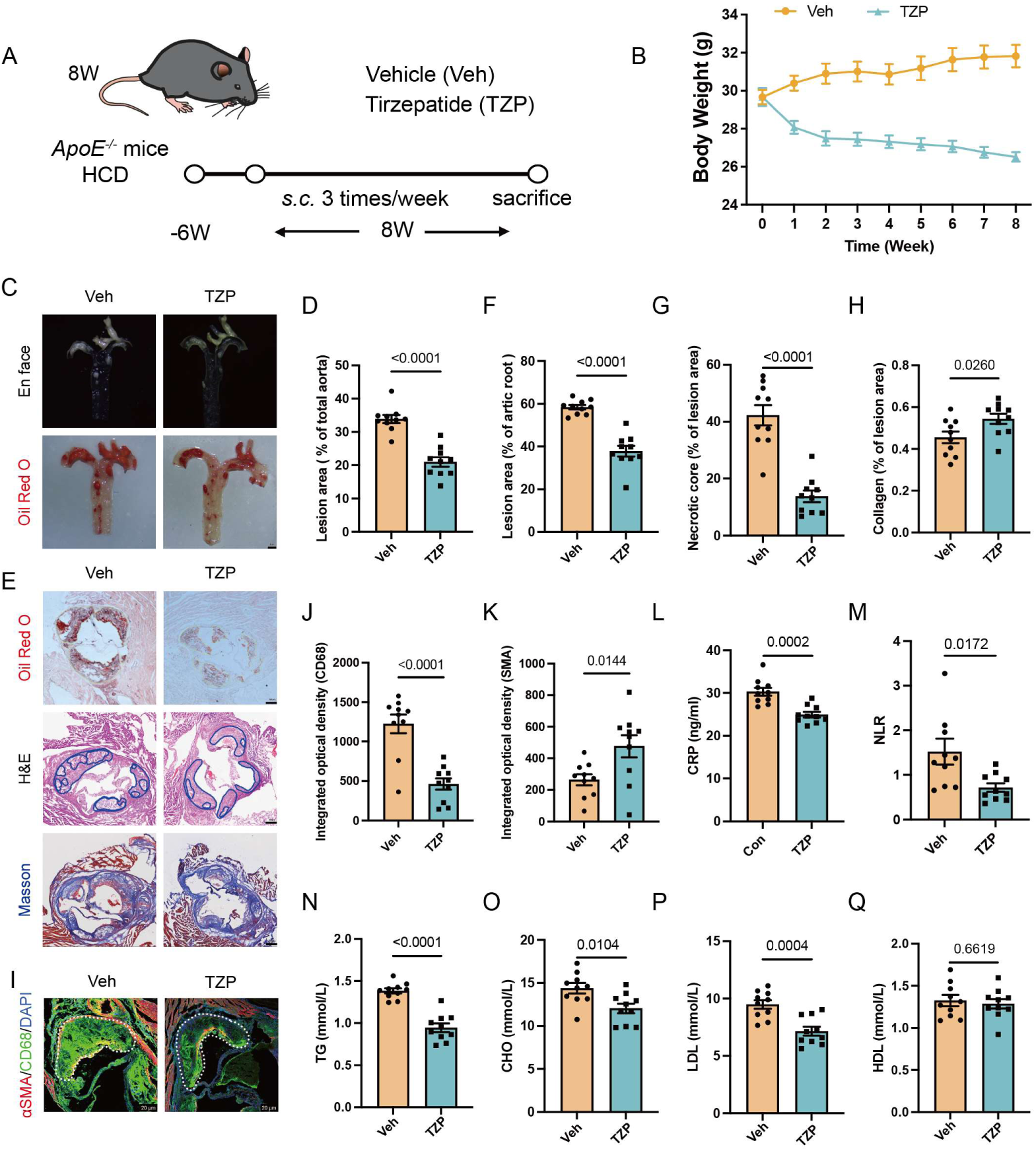
TZP ameliorates atherosclerosis in non-diabetic and non-obese *Apoe^-/-^* mice. **(A)** Schematic overview of the animal experimental design. **(B)** Body weight trajectories of *Apoe^-/-^* mice receiving vehicle (Veh), low-dose tirzepatide (TZP-L), or high-dose tirzepatide (TZP-H). **(C** – **D)** Representative en face Oil Red O staining and quantitative analysis of atherosclerotic lesions isolated from whole aortas. TZP effectively decreased aortic atherosclerotic lesion burden. Scale bar: 1 mm. **(E–H)** Aortic root sections stained with Oil Red O, hematoxylin and eosin (H&E), and Masson trichrome staining with corresponding quantitative analyses. TZP diminished plaque area and necrotic core size while elevating collagen deposition within atherosclerotic lesions. Scale bar: 200 μm. **(I–K)** Representative immunofluorescence images and statistical quantification of aortic root plaques stained for alpha-smooth muscle actin (α-SMA; red) and cluster of differentiation 68 (CD68; green). TZP decreased CD68-positive macrophage infiltration and increased the α-SMA-positive smooth muscle cell area. Scale bar: 20 μm. **(L–M)** Serum C-reactive protein (CRP) concentrations and neutrophil-to-lymphocyte ratio (NLR). TZP intervention rescued aberrant systemic inflammation in *Apoe^-/-^* mice. **(N–Q)** Assessment of serum lipid profiles, including triglyceride (TG), total cholesterol (TC), high-density lipoprotein cholesterol (HDL-C), and low-density lipoprotein cholesterol (LDL-C). High-dose TZP reduced circulating TG levels without altering TC, LDL-C, or HDL-C contents. Data are presented as mean±SEM; n=10 for all assigned groups. Two-way ANOVA coupled with Tukey’s multiple comparisons test was used for body weight analysis in (B). A two-tailed unpaired Student’s t-test was utilized for statistical evaluation across panels (C–K).

Together, these findings extend the anti-atherosclerotic effects of TZP from diabetic atherosclerosis and obese atherosclerosis, to a non-diabetic, non-obese setting, as evidenced by reduced lesion burden, improved lipid metabolism, attenuated inflammation, and favorable changes in plaque composition.

### TZP inhibits atherosclerosis progression through a weight-loss independent manner

To directly assess whether the anti-atherosclerotic effects of TZP are weight-dependent, we performed ANCOVA across three mouse models as described above. In diabetic atherosclerotic mice, the overall ANCOVA model was significant (*F*(2,7)=10.30, *P*=0.00082). A significant main effect of group on plaque area percentage was observed (*F*(1,7)=7.84, *P*=0.02665), whereas body weight was not a significant covariate (*F*(1,7)=0.019, *P*=0.899), indicating that between-group differences were not driven by body weight (**Table S1**). Similarly, in obese mice, the overall model remained significant (*F*(3,14)=11.46, *P*=0.0005), with a significant group effect (*F*(2,14)=6.47, *P*=0.0102) but no effect of body weight (*F*(1,14)=0.80, *P*=0.387), confirming that the observed differences were independent of body weight (**Table S2**). Consistent results were obtained in non-diabetic, non-obese atherosclerotic mice, in which the ANCOVA model was significant (*F*(2,17)=33.03, *P*<0.0001), with a significant group effect (*F*(1,17)=11.41, *P*=0.0036) and no contribution of body weight (F(1,17)=0.41, P=0.531), further supporting weight-independent effects (**Table S3**). Collectively, these findings indicate that the reduction in plaque burden observed after TZP treatment cannot be solely attributed to differences in body weight. Instead, TZP-associated vascular protection persisted after adjustment for body weight across diabetic, obese, and non-diabetic/non-obese atherosclerosis settings, supporting a body weight-independent component of its anti-atherosclerosis action.

To further determine whether the anti-atherosclerotic effects of TZP are independent of body weight, we performed a pair-feeding study in *Apoe^-/-^* mice. After 6 weeks of HCD feeding, mice were assigned to control, TZP-treated, or pair-fed (PF) groups to account for differences in food intake and body weight loss. (**Fig. 4A**).

**Fig. 4.**
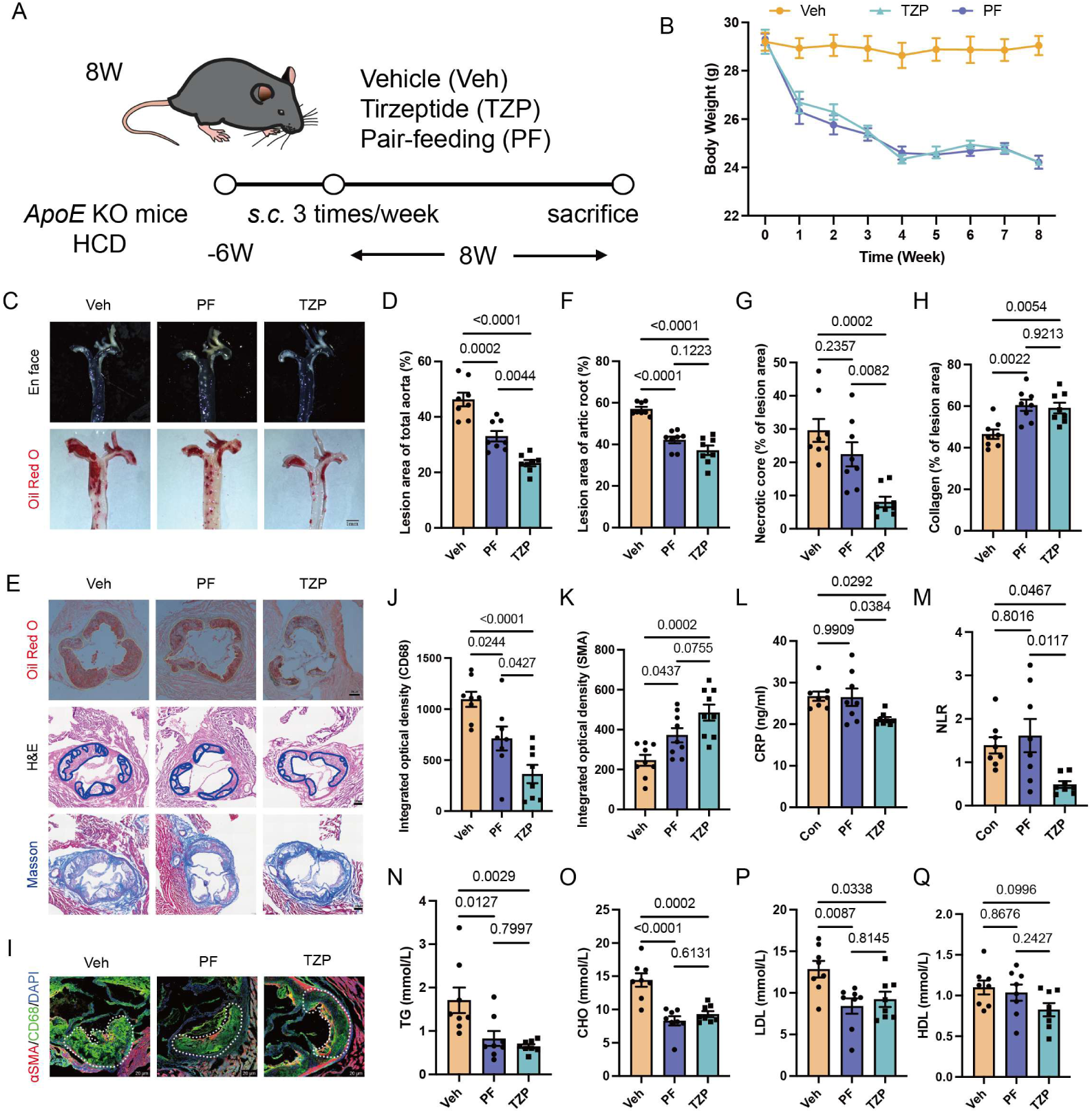
TZP inhibits atherosclerosis progression through a weight loss-independent manner. **(A)** Schematic overview of the overall experimental design. **(B)** Dynamic body weight trajectories of *Apoe^-/-^* mice assigned to vehicle (Veh), pair-fed (PF), and tirzepatide (TZP; 10 nmol/kg) groups. **(C–D)** Representative en face Oil Red O staining and quantitative quantification of atherosclerotic lesions in isolated whole aortas. Scale bar: 1 mm. **(E–H)** Histological evaluation of aortic root sections via Oil Red O, hematoxylin and eosin (H&E), and Masson trichrome staining accompanied by corresponding quantitative analyses. Scale bar: 200 μm. **(I–K)** Representative immunofluorescence images and statistical quantification of aortic root plaques stained for alpha-smooth muscle actin (α-SMA; red) and cluster of differentiation 68 (CD68; green). Scale bar: 20 μm. **(L–M)** Serum C-reactive protein (CRP) levels and neutrophil-to-lymphocyte ratio (NLR), demonstrating that TZP treatment ameliorated systemic inflammation. **(N–Q)** Detection of serum lipid profiles including triglyceride (TG), total cholesterol (TC), low-density lipoprotein cholesterol (LDL-C), and high-density lipoprotein cholesterol (HDL-C). Both PF and TZP interventions decreased serum TG, TC, and LDL-C levels, whereas HDL-C content remained unchanged across all groups. Data are presented as mean±SEM; n=8 for each experimental group. Two-way ANOVA followed by Tukey’s multiple comparisons test was applied for body weight analysis in (B). One-way ANOVA with Bonferroni’s post hoc correction was performed for statistical comparison in panels (C–K).

During the intervention period, body weight declined progressively in both the PF and TZP groups, and the body weight curves of two groups closely overlapped (**Fig. 4B**). This confirmed that the pair-feeding regimen effectively reproduced the weight-loss effects observed in TZP treatment, providing a controlled setting to distinguish weight-dependent from weight-independent vascular effects. Notably, both PF and TZP treatment reduced aortic plaque area compared with control mice. However, TZP treatment produced greater suppression of atherosclerosis than PF group (**Fig. 4C–4D**). Furthermore, Oil Red O staining of the aortic sinus revealed a marked reduction in atherosclerotic plaque area after TZP or PF intervention. Analysis of plaque composition further supported an additional vascular benefit of TZP. Both PF and TZP increased plaque collagen content, with no obvious difference between the two groups. Moreover, necrotic core area was reduced in either TZP or PF groups but was significantly smaller in TZP-treated mice (**Fig. 4E–4H**).

Consistent with these results, CD68⁺ macrophages accumulation was significantly inhibited while α-SMA^+^ smooth muscle cells were increased in plaques from both PF and TZP groups compared to control mice. More importantly, the decrease of macrophage accumulation was more pronounced in TZP-treated mice than that in PF mice, whereas α-SMA^+^ area did not significantly differ between TZP and PF intervention groups (**Fig. 4I–4K**). Inflammatory profiles were further evaluated. Both PF and TZP markedly lowered CRP levels, with a greater decrease in the TZP group. NLR levels was also significantly reduced by TZP but not by PF, and direct comparison confirmed a statistical difference between TZP and PF groups (**Fig. 4L–4M**). Additionally, mice from both TZP and PF groups exhibited lowered serum TG, TC and LDL-C levels, while the level of HDL-C were not significantly altered among three groups (**Fig. 4N–4Q**), indicating that lipid modulation may be dispensable for the additional anti-atherosclerotic mice treated with TZP.

The above data suggest that reduced food intake and body weight loss account for partial vascular benefit observed in TZP intervention group. The greater suppression of plaque burden, necrotic core size, macrophage accumulation, and systemic inflammation in TZP-treated mice supports an additional anti-atherosclerotic mechanism beyond body weight reduction

### scRNA-seq reveals weight-loss independent gene signature in vascular endothelium from atherogenic mice

To further investigate the mechanisms by which TZP attenuates atherosclerosis progression, we performed single-cell RNA sequencing (scRNA-seq) of mouse aortas isolated from experimental mice (**Fig. 5A**). Based on the Z-score normalized expression of canonical cell-type marker genes, we identified nine major cell populations in the aortic tissue, including vascular smooth muscle cells (VSMCs), endothelial cells, fibroblasts, macrophages, T cells, B cells, granulocytes, natural killer cells, and Schwann cells (**Fig. 5B**). Uniform Manifold Approximation and Projection (UMAP) analysis was then used to visualize cellular clustering and distribution in two-dimensional space. Cell composition analysis further demonstrated increased proportion of endothelial cells in the TZP group compared with the vehicle (Veh) group, suggesting partial restoration of vascular endothelial homeostasis and enhanced plaque stability (**Fig. 5C**). Notably, we found that both *Glp1r* and *Gipr* were highly expressed in endothelial cells, suggesting that endothelial cell may serve as a potential TZP-responsive target cell type within atherosclerotic plaque (**Fig. 5D**).

**Fig. 5.**
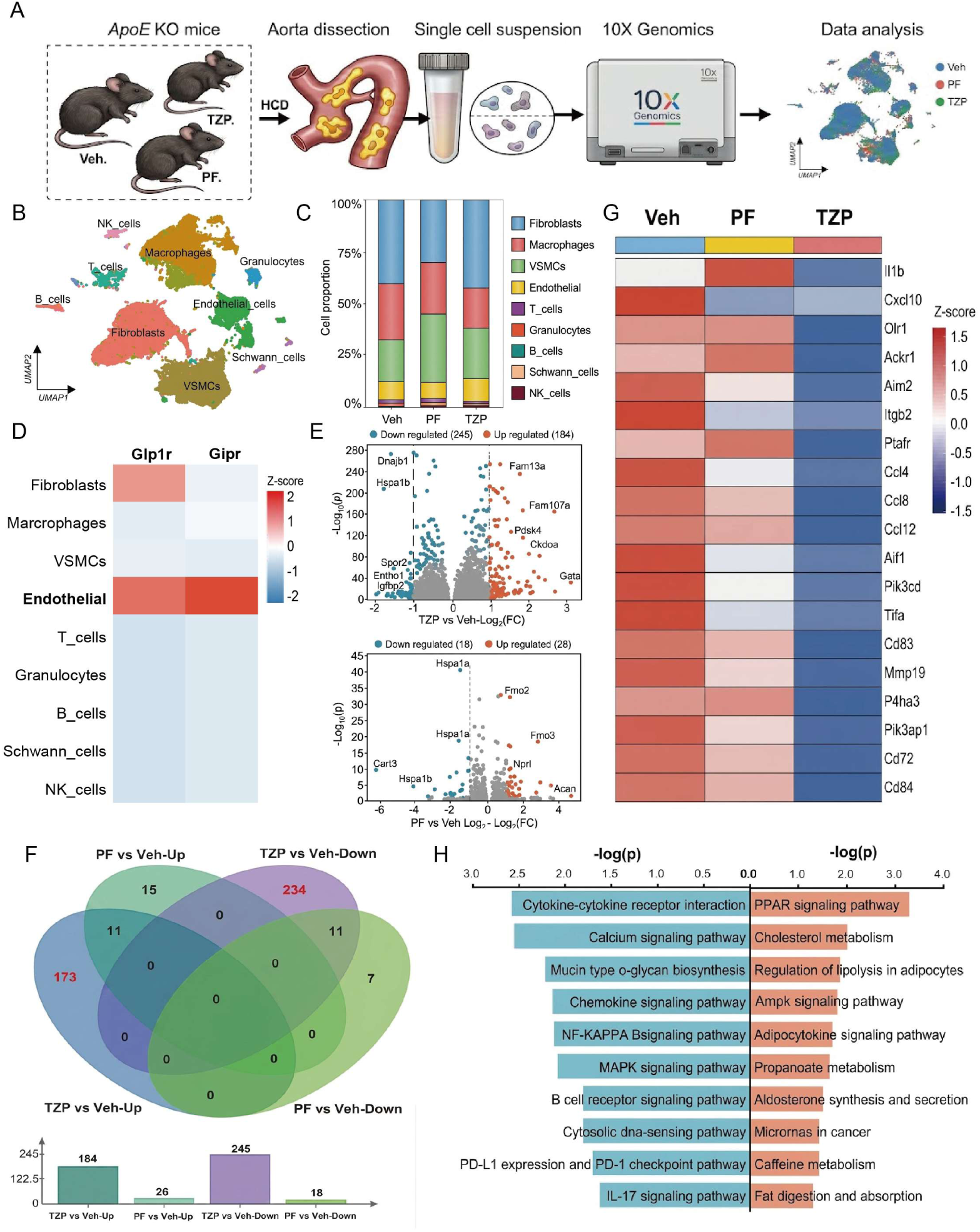
Single-cell RNA-sequencing uncovers weight loss-independent transcriptional signatures in the vascular endothelium of atherogenic mice. **(A)** Schematic workflow of single-cell RNA-sequencing (scRNA-seq) analysis. High-fat diet-fed *Apoe^-/-^* mice received vehicle (Veh), pair-fed (PF), or tirzepatide (TZP) treatment. Aortic tissues harvested from 3 mice per group were pooled and subjected to 10× Genomics scRNA-seq and subsequent bioinformatic analyses. **(B)** Uniform manifold approximation and projection (UMAP) visualization illustrating distinct cell clusters isolated from mouse aortic single-cell suspensions, color-coded according to annotated cell types. **(C)** Bar graph showing the proportional distribution of aortic cellular components across the Veh, PF, and TZP experimental groups. **(D)** Heatmap illustrating the transcript abundance of glucagon-like peptide 1 receptor (*Glp1r*) and gastric inhibitory polypeptide receptor (*Gipr*) across all identified aortic cell subtypes. **(E)** Volcano plots summarizing differentially expressed genes (DEGs) within endothelial cells. Comparisons were performed for TZP versus Veh (upper panel) and PF versus Veh (lower panel); representative DEGs were manually annotated. **(F)** Venn diagram depicting the overlap of DEGs derived from endothelial cells among indicated comparative groups. **(G)** Heatmap profiling the expression patterns of inflammation-associated genes that were significantly suppressed in endothelial cells after experimental interventions. **(H)** Kyoto Encyclopedia of Genes and Genomes (KEGG) pathway enrichment analysis of screened DEGs. Enriched upregulated pathways are highlighted in orange, and downregulated pathways are highlighted in blue.

To further investigate endothelial-specific transcriptional changes, we performed differential gene expression in endothelial cells across different groups (**Fig. 5E**). After adjusting for body weight-related effects, comparison with the control group identified 173 upregulated and 234 downregulated differentially expressed genes (DEGs) in endothelial cells following TZP administration (**Fig. 5F**). Consistent with the ameliorated inflammatory phenotype observed in vivo, z-score normalized heatmap visualization of representative pro-inflammatory DEGs revealed that these genes were predominantly enriched in the Veh group, exhibited intermediate expression levels in the PF group, and were markedly repressed upon TZP treatment (**Fig. 5G**).

To further elucidate the functions of these DEGs, we performed a Kyoto Encyclopedia of Genes and Genomes (KEGG) analysis on both the upregulated and downregulated sets. The top 10 upregulated and downregulated pathways identified through KEGG pathway enrichment analysis are illustrated in the accompanying figure (**Fig. 5H**). KEGG pathway enrichment analysis revealed that TZP intervention significantly upregulated signaling pathways associated with metabolic protection—including the adenosine monophosphate-activated protein kinase (AMPK) signaling pathway, Peroxisome proliferator-activated receptor (PPAR) signaling pathway, cholesterol metabolism, and adipokine signaling pathway—while simultaneously significantly inhibiting pro-inflammatory pathways such as the Nuclear factor kappa-light-chain-enhancer of activated B cells (NF-κB) signaling pathway, chemokine signaling pathway, and cytokine-cytokine receptor interaction. These results align closely with the anti-inflammatory effects of TZP in the vasculature.

### TZP activates eNOS and reduces inflammation in normal C57BL/6J mice challenged with TNF-α

To further examine the anti-inflammatory role of TZP *in vivo*, we established an acute TNF-α-induced inflammation model in mice. Eight-week-old mice fed with normal chow diet were randomly assigned to control, TNF-α, or TZP plus TNF-α groups, with no significant differences in baseline body weight and fasting glucose among three groups (**Fig. 6A–6C**). Mice were pretreated with vehicle or TZP, before intraperitoneal injection of TNF-α to elicit acute inflammation. Six hours after TNF-α challenge, mice were euthanized and tissues as well as serum were collected for analysis.

**Fig. 6.**
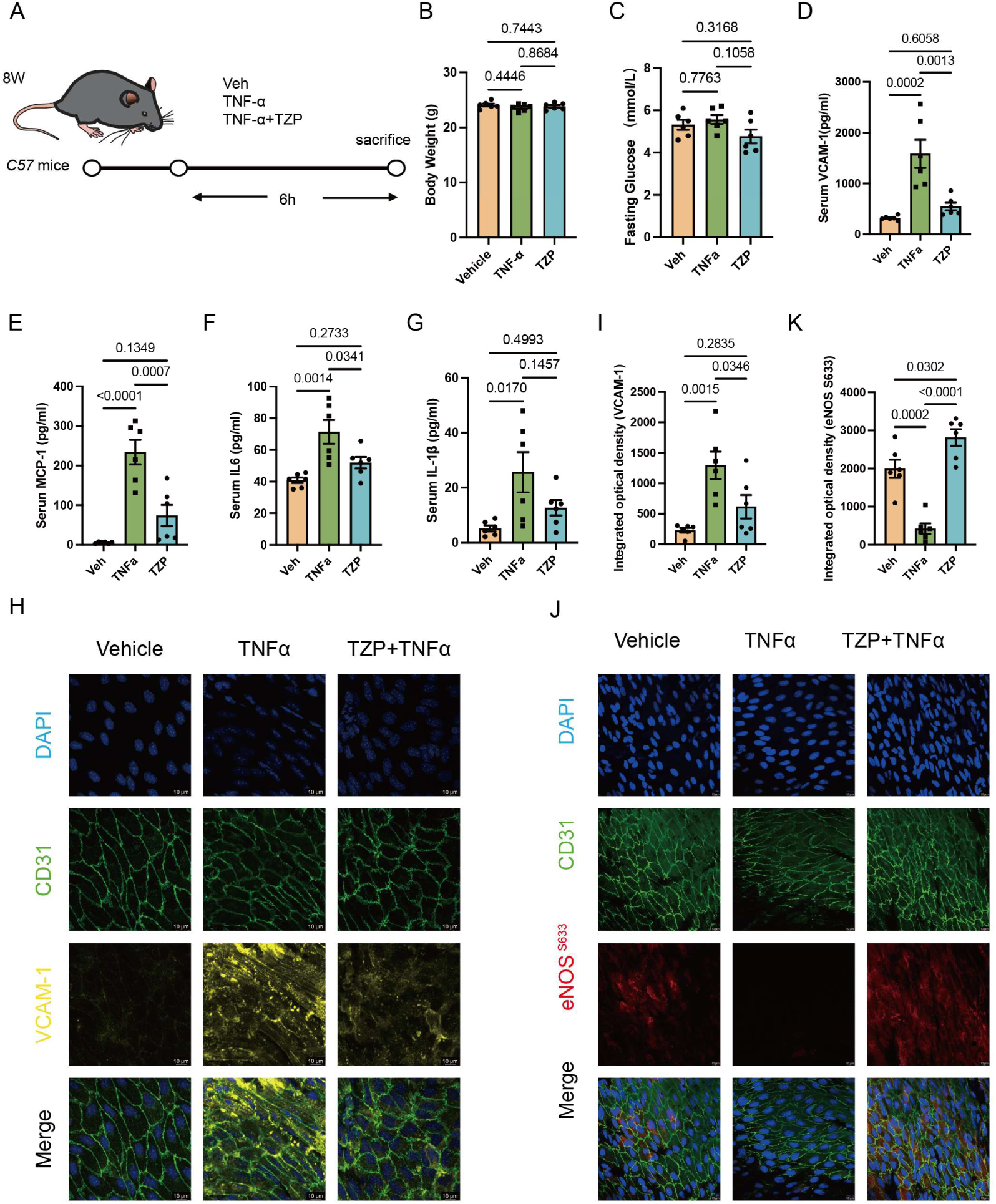
TZP activates eNOS signaling and alleviates vascular inflammation in TNF-α-stimulated C57BL/6J mice. **(A)** Schematic diagram illustrating the in vivo experimental design. C57BL/6J mice were pretreated with vehicle (Veh) or tirzepatide (TZP), followed by intraperitoneal administration of tumor necrosis factor-α (TNF-α) or vehicle control. All mice were euthanized 6 hours after modeling. **(B–C)** The alterations in body weight and fasting blood glucose among different treatment groups. **(D–G)** Quantification of serum inflammatory biomarkers, including vascular cell adhesion molecule-1 (VCAM-1), monocyte chemoattractant protein-1 (MCP-1), interleukin-6 (IL-6), and interleukin-1β (IL-1β). **(H)** Representative en face immunofluorescence images of aortic vascular cell adhesion molecule-1 (VCAM-1) expression. Scale bar: 50 μm. **(I)** Quantitative analysis of VCAM-1 immunofluorescence intensity corresponding to panel (H). **(J)** Representative en face immunofluorescence staining for phosphorylated endothelial nitric oxide synthase (p-eNOS; Ser633) in isolated aortas. Scale bar: 50 μm. **(K)** Statistical quantification of phosphorylated eNOS (Ser633) immunofluorescence signals shown in panel (J). **Data are presented as mean±SEM**; n=6 for each group. One-way ANOVA with Bonferroni’s multiple comparisons test was utilized for statistical analyses for panels (B–G), (I), and (K).

Multiplex cytokine profiling showed that TNF-α significantly upregulated the levels of circulating inflammatory factors, including vascular cell adhesion molecule-1 (VCAM-1), Monocyte chemoattractant protein-1 (MCP-1), interleukin-6 (IL-6), and interleukin-1β (IL-1β), while treatment with TZP statistically reduced the serum VCAM-1 and MCP-1 levels (**Fig. 6D-6G**). Consistently, *en face* immunofluorescence staining of arterial vessels revealed decreased endothelial VCAM-1 expression in TZP-pretreated mice than that in mice receiving TNF-α alone (**Fig. 6H–6I**). These findings demonstrate that TZP suppresses both systemic and vascular endothelial inflammation in response to TNF-α challenge.

Given that TNF-α-mediated vascular inflammation is closely linked to endothelial dysfunction, we next investigated whether TZP preserves endothelial protective signaling under inflammatory stress. Nitric oxide (NO) produced by endothelial nitric oxide synthase (eNOS) plays a central role in regulating endothelial homeostasis. Phosphorylation of eNOS at serine residue 633 (Ser633, corresponding to Ser635 in mouse eNOS) has been suggested to be a pivotal post-translational modification that maintains endothelial function, enhances NO bioavailability, and inhibits endothelial inflammatory activation. Inflammatory stress such as TNF-α exposure robustly impairs eNOS Ser633 phosphorylation, thereby aggravating endothelial injury and atherosclerotic progression^22,23^. Immunofluorescence staining revealed TNF-α stimulation reduced endothelial eNOS Ser633 phosphorylation, which was significantly restored by TZP treatment (**Fig. 6J–6K**).

### TZP inhibits endothelial inflammation

We demonstrated that TZP attenuates systemic and vascular inflammation, which may contribute to its anti-atherosclerotic effects. Consistent with our single-cell sequencing data, previous studies have reported that GLP-1R and GIPR are highly expressed in vascular endothelial cells^24–27^. We first assessed the cytotoxicity of TZP by measuring cell viability in human umbilical vein endothelial cells (HUVECs) (**Fig. 7A**). Considering the pivotal role of endothelial activation in leukocyte recruitment, vascular inflammation, and atherosclerosis progression, we next explored the anti-inflammatory effects of TZP. We established an inflammatory model by stimulating HUVECs with TNF-α, and subsequently investigated whether TZP treatment could effectively attenuate endothelial cell inflammation. We observed that TZP treatment significantly reversed TNF-α induced endothelial inflammation (**Fig. 7B-7C**). Furthermore, western blot analysis revealed that TZP reversed the upregulation of the adhesion molecules VCAM-1 and ICAM-1 in endothelial cells in a dose-dependent manner (**Fig. 7D**). The anti-inflammatory effect of TZP was also recapitulated in human aortic endothelial cells (HAECs)(**Fig. S4F-4G**).

**Fig. 7.**
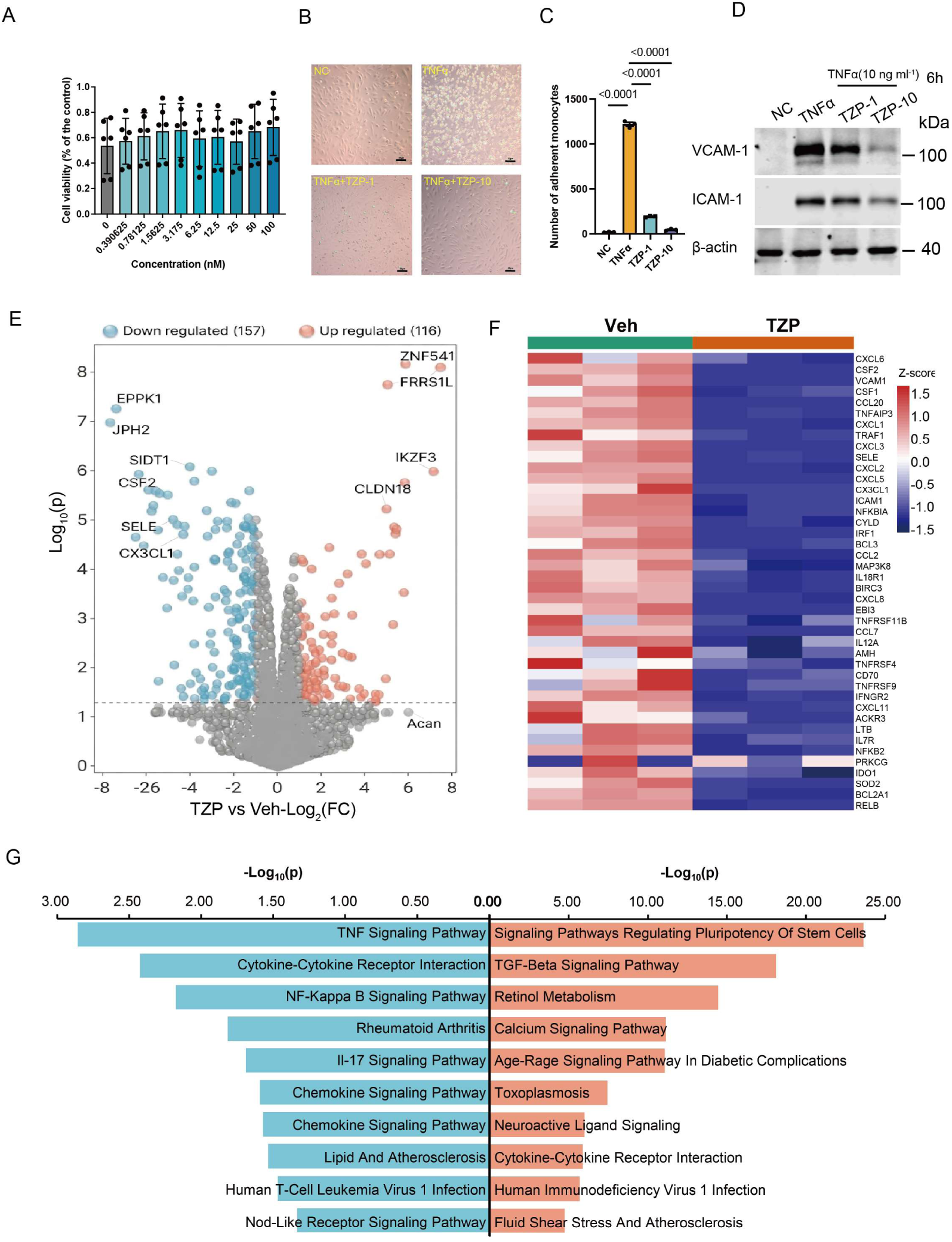
TZP suppresses inflammatory responses in human vascular endothelial cells. **(A)** Cell viability assessment of human umbilical vein endothelial cells (HUVECs) incubated with gradient concentrations of TZP for 24 hours via cell counting kit-8 (CCK-8) assay (n=6). **(B)** Representative fluorescence images showing the adhesion of calcein-AM-labeled THP-1 monocytes to tumor necrosis factor-α (TNF-α)-stimulated HUVECs. TZP treatment markedly inhibited monocyte-endothelial adhesion (n=3). Scale bar: 50 μm. **(C)** Quantitative statistics of adhered THP-1 monocytes corresponding to panel (B). **(D)** Western blot detection and quantitative analysis of vascular cell adhesion molecule-1 (VCAM-1) and intercellular adhesion molecule-1 (ICAM-1) protein expression in HUVECs (n=3). **(E)** Volcano plot illustrating differentially expressed genes (DEGs) between vehicle (Veh)-treated and TZP-treated HUVECs for transcriptomic profiling (n=3). **(F)** Heatmap displaying the expression patterns of significantly downregulated inflammation-related DEGs in TZP-intervened HUVECs. **Data are presented as mean±SEM**. One-way ANOVA followed by Bonferroni’s multiple comparisons test was applied for statistical analyses across panels (B–F).

To investigate the global transcriptional effects of TZP on TNFα-stimulated endothelial cells, we performed RNA-sequencing analysis in HUVECs. The volcano plot (**Fig. 7E**) shows the distribution of differentially expressed genes (DEGs) between TZP-treated and vehicle control groups. In total, 116 genes were upregulated in response to TZP treatment, while 157 genes were significantly downregulated. Notably, many pro-inflammatory genes, including chemokines C-X-C motif chemokine ligand (CXCL) 1, CXCL2, CXCL3, C-C motif chemokine ligand (CCL) 2, and colony-stimulating factor 2 (CSF2), were among the most significantly downregulated transcripts, whereas genes related to cell junction maintenance and transcriptional regulation, such as claudin 18 (CLDN18), were upregulated. Consistent with the volcano plot, the heatmap analysis (**Fig. 7F**) further revealed that TZP treatment broadly suppressed the expression of a plethora of TNFα-induced inflammatory and adhesion-related genes, including VCAM1, ICAM1, Interleukin 6 (IL6), and multiple chemokines, compared to the vehicle control.

To identify the biological pathways modulated by TZP, we performed KEGG pathway enrichment analysis (**Fig. 7G**). Genes downregulated by TZP were significantly enriched in pathways associated with inflammatory responses and atherosclerosis, including the TNF signaling pathway, cytokine-cytokine receptor interaction, NF-κB signaling pathway, chemokine signaling pathway, IL-17 signaling pathway, and lipid and atherosclerosis signaling. In contrast, upregulated genes were enriched in pathways related to cellular homeostasis and protective functions, such as the TGF-β signaling pathway and calcium signaling pathway.

Collectively, these data suggest that Tirzepatide can effectively attenuate inflammation within endothelial cells.

### Activation of cAMP/PKA/eNOS pathway is involved in the anti-inflammatory effects of TZP in endothelial cells

Tirzepatide is a dual GLP-1R/GIPR agonist. To determine whether it’s mechanism of action is dependent on its canonical receptors, we utilized siRNA in HUVECs to selectively knock down GLP-1R, GIPR, or both receptors simultaneously in HUVECs (knockdown validation in **Fig. S5A-4B**). The results demonstrated that knockdown of GLP-1R partially reversed the anti-inflammatory effects of TZP, whereas simultaneous knockdown of both GLP-1R and GIPR completely abrogated the drug’s anti-inflammatory activity, suggesting that the anti-inflammatory effects of TZP are mediated through GLP-1R and GIPR (**Fig. 8A–8C**). GLP-1R and GIPR both belong to the G protein-coupled receptor (GPCR) family. The Gαs/cyclic adenosine monophosphate (cAMP) signaling pathway is a classic downstream cascade reaction activated by GPCRs^28,29^.

**Fig. 8.**
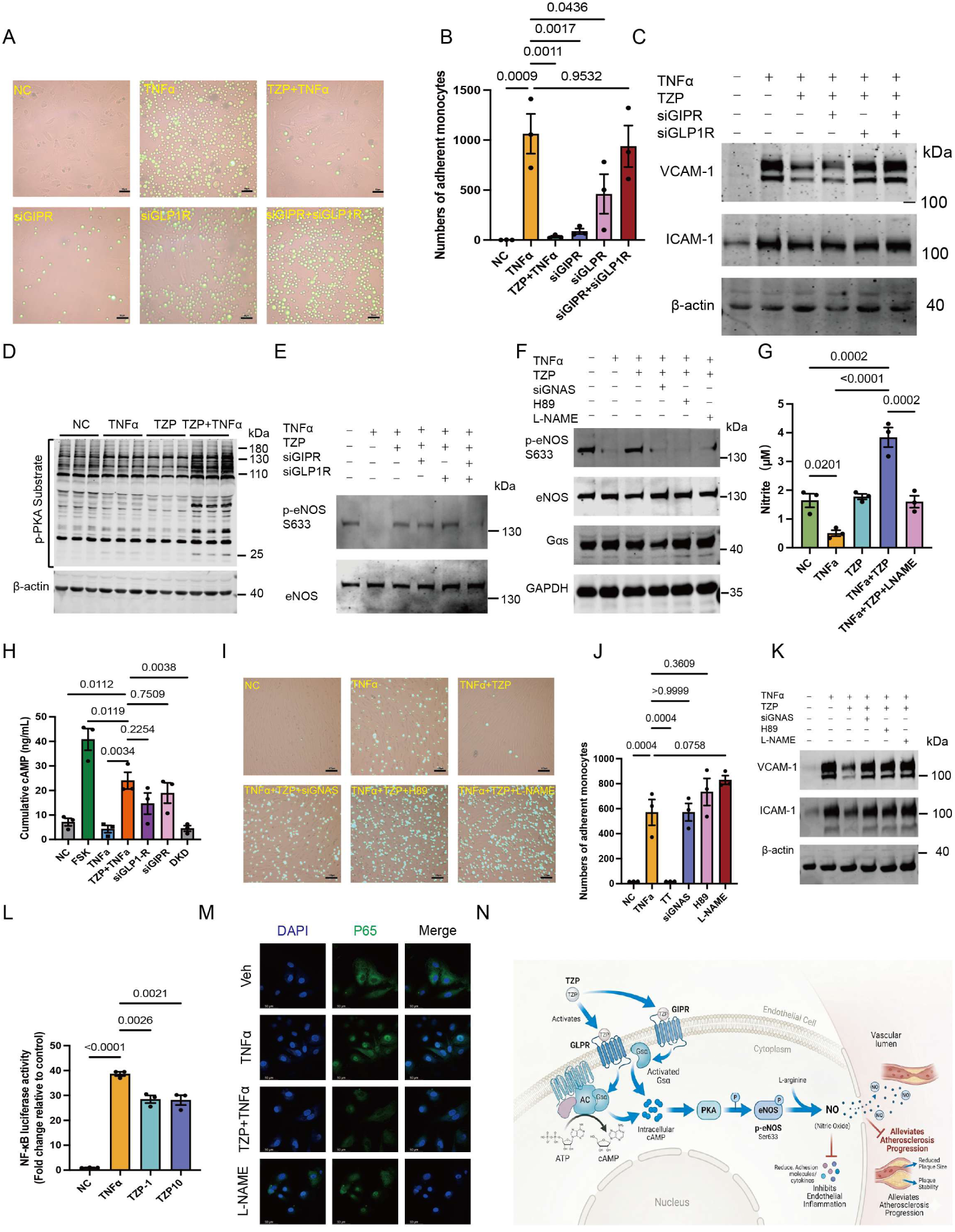
Tirzepatide blunts endothelial inflammation via activation of the cyclic adenosine monophosphate (cAMP)/protein kinase A (PKA)/endothelial nitric oxide synthase (eNOS) signaling cascade. **(A)** Representative microscopic images showing the effects of small interfering RNA (siRNA)-mediated knockdown of glucagon-like peptide-1 receptor (GLP-1R), gastric inhibitory polypeptide receptor (GIPR), or dual receptor knockdown on TZP-suppressed THP-1 monocyte adhesion to HUVECs. Scale bar: 50 μm. **(B)** Quantitative analysis of adhered THP-1 monocytes corresponding to the experimental groups described in (A). **(C)** Immunoblot analysis of vascular cell adhesion molecule-1 (VCAM-1) and intercellular adhesion molecule-1 (ICAM-1) expression following GLP-1R and/or GIPR silencing in HUVECs. **(D)** Immunoblot detection of phosphorylated PKA substrates in tumor necrosis factor-α (TNF-α)-challenged HUVECs with or without TZP intervention. **(E)** Western blot analysis to evaluate the regulatory roles of GLP-1R and GIPR in TZP-triggered endothelial nitric oxide synthase (eNOS; Ser633) phosphorylation using targeted siRNA interference. **(F)** Evaluation of TZP-induced eNOS (Ser633) phosphorylation upon the inhibition of stimulatory G protein (GNAS) via siGNAS, PKA blockade using H89, and eNOS inhibition using Nω-nitro-L-arginine methyl ester (L-NAME). **(G)** Quantification of nitric oxide (NO) production in the cell culture supernatant collected from TZP-treated HUVECs. **(H)** Measurement of intracellular cAMP accumulation in HUVECs after individual or combined knockdown of GLP-1R and GIPR (n=3). **(I)** Representative adhesion images illustrating the impacts of siGNAS, H89, and L-NAME on TZP-mediated inhibition of THP-1 monocyte adhesion to endothelial cells. Scale bar: 100 μm. **(J)** Statistical quantification of adhered THP-1 monocytes corresponding to panel (I). **(K)** Dual-luciferase reporter assay for the evaluation of nuclear factor κB (NF-κB) transcriptional DNA-binding activity under indicated treatments. **(L)** Immunofluorescence staining to characterize the nuclear translocation of NF-κB p65 subunit in response to different experimental interventions. **(M–N)** Schematic working models summarizing the underlying molecular mechanisms. TZP ameliorates endothelial inflammatory injury and retards atherosclerotic progression through GLP-1R/GIPR-dependent Gs/cAMP/PKA/eNOS signaling, thereby restraining NF-κB-mediated proinflammatory activation. Data are presented as mean±SEM; n=3 for all experimental groups. One-way ANOVA with Tukey’s multiple comparisons test was performed for statistical analyses of panels (B), (H), (J), and (L).

By examining phosphorylated protein kinase A (PKA) target proteins—the key downstream effector molecules of cAMP—we confirmed that, under inflammatory conditions, PKA signaling in TZP-treated endothelial cells was activated (**Fig. 8D**). Given that PKA can directly phosphorylate eNOS at serine residue 633 (Ser633)^30^, we first characterized the temporal phosphorylation pattern of this functionally critical eNOS site upon TNF-α stimulation alone: Ser633 represents the canonical PKA-dependent activating residue. In HUVECs, TNF-α stimulation exerted divergent time-dependent effects: p-eNOS Ser633 displayed a mild gradual increase across 0–120 min (representative blots shown in **Fig.S5D**). We further investigated whether TZP treatment could upregulate p-eNOS Ser633 activation. In HUVECs, we observed that the inhibitory effect of TNF-α on eNOS Ser633 phosphorylation could be reversed by TZP treatment, and this effect vanished after knocking down GLP1R and GIPR (**Fig. 8E**). To determine whether TZP activation of eNOS is mediated through the cyclic adenosine monophosphate (cAMP)/protein kinase A (PKA) pathway, we selectively inhibited different mediators in this pathway using siGNAS (knockdown validation in **Fig.S5 C**), H89 (a PKA inhibitor), and Nω-nitro-L-arginine methyl ester (L-NAME) (a inhibitor of nitric oxide synthase), respectively, and then monitored the phosphorylation status of eNOS at Ser633 (**Fig. 8F**). Simultaneously, we assessed cAMP and NO levels in HUVECs by measuring the content of cAMP and nitrate content (**Fig. 8G–8H**). Under inflammatory conditions, TZP significantly increased the NO content in the supernatant—an effect that could be inhibited by L-NAME, an inhibitor of eNOS. Correspondingly, further blockage of the corresponding targets in the cAMP/PKA/eNOS pathway reversed the TZP-induced inhibition on monocyte adhesion and the expression of inflammatory markers ICAM-1 and VCAM-1 (**Fig. 8I** – **8K**).

Integrating the results from the aforementioned scRNA-seq and transcriptome profiling, the enriched pathways suggested that the NF-κB pathway is inhibited by TZP. To validate whether the NF-κB pathway in endothelial cells mediates the anti-inflammatory effects downstream of eNOS, we employed a dual-luciferase reporter assay system. The results demonstrated that TZP effectively inhibits TNF-α-induced DNA binding activity of NF-κB p65 (**Fig. 8L**). Immunofluorescence staining of p65, further corroborate that TZP effectively suppresses the nuclear translocation of p65, which can this effect could be reversed by L-NAME (**Fig. 8M**).

Collectively, these findings indicate that TZP treatment alleviates endothelial cell inflammation by activating the cAMP/PKA/eNOS pathway. This activation inhibits the downstream NF-κB signaling cascade, thereby suppressing the expression of pro-inflammatory mediators and restoring endothelial resilience (**Fig. 8N**), which may be translatable into its atheroprotective effects *in vivo*.

## Discussion

In the present study, we demonstrated that tirzepatide (TZP) exerts anti-atherosclerotic effects across three distinct mouse models, recapitulating the atherosclerotic phenotypes associated with diabetes, obesity, and non-diabetic/non-obese metabolic status, respectively. Crucially, our findings resolve a long-standing clinical conundrum: whether the cardiovascular benefits of this dual GLP-1R/GIPR agonist stem merely from systemic metabolic amelioration or from a direct vascular protective mechanism^31,32^. While nearly all participants in previous cardiovascular outcome trials presented confounding concomitant metabolic disorders, our integration of systematic phenotyping, and the use of ANCOVA, non-diabetic/non-obese model, an acute inflammation model as well as pair feeding experiments firmly establishes that the atheroprotective action of TZP is, at least in part, independent of body weight loss and metabolic improvement.

Mechanistically, we delineated that TZP elicits direct anti-inflammatory effects in vascular endothelial cells via activation of the cAMP/PKA/eNOS signaling axis, which subsequently suppresses downstream NF-κB-mediated inflammatory cascades. Our work extends significantly beyond these observations. By uncoupling direct endothelial protection from secondary metabolic consequences, this study provides direct evidence validating the intrinsic vascular benefits of dual receptor agonism, offering deeper insights into its clinical cardiovascular actions.

The pharmacological actions and mechanisms of TZP in vascular diseases have aroused intensive research interest. For example, TZP has been recently reported to mitigate thoracic aortic aneurysm and dissection by alleviating VSMC phenotypic switch and reducing vascular inflammation^33^. Furthermore, two recent preclinical studies on TZP related to atherosclerosis have yielded divergent conclusions. One of which showed that TZP can downregulate inflammatory factors such as MCP-1, VCAM-1, and IL-6, inhibiting aortic plaque progression in early stage diabetic *Apoe^-/-^*mice; however, TZP had no significant protective effect on late-stage diabetic mice or non-diabetic mice^34^. Another concurrent study confirmed that TZP has vasculoprotective effect in *ApoE^⁻/⁻^* mice, and in vitro experiments verified its ability to regulate macrophage polarization and reshape M1/M2 homeostasis^35^. However, the above studies do not address whether the anti-atherosclerotic effects of TZP are derived from its weight-loss effects or direct effects on the vasculature. Our independent study supports these findings, confirming that TZP exerts vasculoprotective effect in both diabetic/obese and non-diabetic/non-obese mouse models. Through strict dietary control, we eliminated confounding factors such as TZP’s appetite suppression and weight reduction, clarifying that its anti-atherosclerotic effect is independent of its metabolic disorder improvement mechanism. These findings could broaden the potential applicability of TZP by addressing its direct vasculoprotective effects and expand its therapeutic value in cardiovascular diseases accompanying metabolic disorders.

Notably, in the obesity-associated atherosclerotic model, low-dose TZP markedly lowered body weight and food intake but failed to achieve a comparable reduction in atherosclerotic lesion formation. This clear phenotypic dissociation indicates that weight loss *per se* is insufficient to account for TZP-mediated atheroprotection, strongly implying an inherent weight-independent mechanism underlying its anti-atherosclerotic action. In line with these preclinical phenotypic findings, a rigorously designed real-world longitudinal cohort study adopting the target trial emulation framework leveraging U.S. Veterans Affairs administrative healthcare data documented progressively elevated MACE risk following GLP-1RA discontinuation; this adverse risk gradient persisted after comprehensive multivariable adjustment for baseline body weight and multiple metabolic confounders, furnishing high-quality population-level evidence supporting weight-dissociated cardioprotection across the GLP-1RA drug class^36^. To rigorously disentangle the confounding influences of body weight and intrinsic vascular protection, we employed paired dietary control experiments. Our results further confirmed that the anti-atherosclerotic efficacy of TZP is preserved after eliminating body weight interference, definitively establishing that TZP protects against atherosclerosis in a weight-independent manner.

Growing clinical evidence underscores the central role of vascular inflammation in atherosclerotic cardiovascular disease. The approval of colchicine by FDA for secondary cardiovascular prevention further highlights residual inflammation as a pivotal therapeutic target in atherosclerotic patients^37,38^. Elevated systemic inflammation, reflected by biomarkers such as CRP, IL-6 and NLR are tightly associated with cardiovascular events^39–41^, raising the possibility of targeting inflammation for risk prediction and prevention of ASCVD in high-risk individuals. Existing clinical data have shown that TZP lowers circulating CRP levels and mitigates inflammatory responses in patients with diabetes and obesity^42,43^. Consistent with these clinical observations, our *in vivo* data across multiple atherosclerotic mouse models, covering both diabetic/obese and non-diabetic lean metabolic backgrounds, demonstrate that TZP profoundly suppresses systemic inflammation and significantly decreases serum CRP and NLR levels. Most importantly, stringent paired-feeding assays normalized body weight across groups and completely eliminated weight-dependent metabolic confounders, further confirming that TZP-mediated anti-inflammatory effects are dissociable from its weight-lowering actions. This scenario is reminiscent of the anti-inflammatory actions of GLP-1 based therapies beyond metabolic benefits^44^.

Uniquely, our single-cell RNA sequencing (scRNA-seq) profiling of aortic tissues from paired-fed mice unveiled the cellular topography underlying this direct vascular action, revealing that *Glp1r* and *Gipr* transcripts were predominantly localized and most highly enriched within the vascular endothelial cell cluster, which is consistent with literature reporting the presence of *Glp1r* in lymphatic endothelium^45^, retinal endothelium^46^, hepatic endothelium^47^ and aortic endothelium^48^. Our study echoes the vascular endothelium as the putative primary target of TZP. To mechanistically recapitulate this receptor-enriched microenvironment, we challenged human endothelial cells with TNF-α *in vitro*. Our transcriptomic analysis revealed that TZP directly prevented endothelial activation, robustly suppressing core gene networks associated with the TNF-α cascade, NF-κB signaling, chemokine pathways, and cytokine-receptor interactions. These cell-intrinsic transcriptomic alterations provide a definitive molecular explanation for the *in vivo* atheroprotection observed in our models, reinforcing the paradigm that TZP directly fortifies the vascular wall against inflammatory insults through a receptor-dependent, endothelium-specific axis across diverse metabolic backgrounds.

Preclinical and clinical studies have shown that GLP-1R activation can slow the progression of atherosclerosis, regardless of hyperglycemia and obesity status^49,50^. In contrast, the specific localization of GIPR in vascular endothelium and the role of GIPR activation in atherosclerosis remains unclear. Although GIPR activation may also mediate weight-independent inflammation-modulatory effects, strong clinical evidence supporting this view is currently lacking^51^. Our *in vitro* experiments using HUVECs revealed that individual knockdown of either GLP-1R or GIPR failed to abolish TZP-mediated restoration of eNOS Ser633 phosphorylation suppressed by TNF α. Prior work has established that dual GLP-1R/GIPR agonists facilitate spatial clustering of membrane-localized GLP-1R and GIPR to form heteromeric receptor nanodomains; such conformational rearrangement synergistically amplifies downstream stimulatory G protein (Gs)/cAMP signaling, accounting for the superior insulinotropic potency of dual agonists relative to selective GLP-1R monoagonists in pancreatic β-cells^52^. In endothelial cells depleted of a single receptor subtype, residual intact receptors remain competent to engage TZP and independently trigger cAMP signaling cascades. Given the existence of a defined activation threshold for the cAMP/PKA axis, ligand stimulation of either receptor alone suffices to drive sufficient cAMP accumulation to cross this threshold and subsequently promote eNOS phosphorylation. This mechanistic framework therefore explains why single-gene knockdown of GLP-1R or GIPR produced no obvious impairment in TZP-induced eNOS activation. This receptor-specific hierarchy potentially provides a mechanistic rationale for the recent SURPASS-CVOT findings; despite achieving superior metabolic and weight-loss outcomes, TZP did not demonstrate an overwhelming superiority over dulaglutide (a selective GLP-1RA) regarding hard cardiovascular endpoints.^10^. The relative contribution of GIPR activation to cardiovascular actions of TZP warrants further exploration.

Several limitations of this study should be acknowledged. First, although we detected divergent reliance on GLP-1R versus GIPR for TZP’s anti-inflammatory actions via cellular adhesion assays and immunoblotting, the precise molecular determinants underlying such receptor-specific signaling disparities remain unresolved owing to current technical constraints. Future validation employing more clinically relevant atherosclerotic animal models or human vascular organoids is therefore warranted^53–55^. For example, conditional deletion of GLP-1R and/or GIPR in vascular organoid endothelial cells, combined with fluorescently labeled TZP tracing, could help delineate receptor engagement dynamics, endocytic trafficking, downstream cAMP signaling, and potential anti-inflammatory signaling bias. Second, constrained by limited sample size, we could not simultaneously adjust for multiple metabolic covariates in statistical analyses. Third, our mechanistic focus was primarily confined to endothelial cAMP/PKA/eNOS/NF-κB signaling; we cannot exclude potential contributions of TZP to functions of vascular smooth muscle cells and myeloid cells (despite *Glp1r* expression in myeloid and hematopoietic cells is almost undetectable^45,56^), nor indirect protective effects mediated by inter-organ crosstalk involving the liver and adipose tissue cannot be fully ruled out. In addition, endothelial cell-specific GLP-1R/GIPR double knockout mice are required to definitively dissect the cell-autonomous role of endothelial GLP-1R and GIPR in mediating TZP’s vasculoprotective actions. Also, atherosclerosis studies in *Glp1r*^Wnt1-/-^ mice (which is resistant to GLP-1RA-induced weight loss^47^) are needed to verify the contribution of food intake suppression to TZP mediated atheroprotection complementary to pair feeding experiments. Lastly, this study only used male mice for animal experiments and did not include female mice. Due to the fluctuations in the menstrual cycle, dynamic changes in endogenous hormones, and individual differences in hormone rhythms in female mice, subsequent animal experiments with females are warranted.

In conclusion, the present study establishes the novel weight-loss independent, anti-atherosclerotic effects of TZP in preclinical mouse models. Mechanistically, TZP restrains endothelial cell inflammation by activating cAMP/PKA/eNOS pathway to suppress NF-κB activation, thereby underlying its atherosclerotic effects across diabetic, obese, and non-diabetic/non-obese conditions. Our findings support the potential clinical utility of TZP in individuals with high cardiovascular risk with or without overt metabolic disorders.

## Acknowledgements

This study was supported by grants from the National Natural Science Foundation of China (grant nos. 82370444 and 12411530127). This study was also supported by USTC research funds of the Double First-Class Initiative (YD9110002089) and the research funds of the Centre for Leading Medicine and Advanced Technologies of IHM (2025IHM01040).

## Author Contributions

S.X., J.W. and Z.W. conceptualized the experiments. S.C., S.W., W.W., T.T., H.L., M.S., F.Z., Y.Y., M.C. and L.J., performed the experiments. P.C.E., B.C.B., S.O. and Y.C. interpreted the data and revised the manuscript for important intellectual content. S.C. and S.X. wrote and edited the final manuscript, with input and approval from all authors.

## Supplemental Material

Materials and Methods

Supplemental Figures S1-S5

Supplemental Tables S1-S4

References 56-62

Major Resource Table

## Nonstandard Abbreviations and Acronyms

GIPR: Glucose-Dependent Insulinotropic Polypeptide Receptor
GLP-1R: Glucagon-Like Peptide-1 Receptor
TZP: Tirzepatide
NLR: Neutrophil-to-Lymphocyte Ratio
VCAM-1: Vascular Cell Adhesion Molecule-1
ICAM-1: Intercellular Adhesion Molecule-1
PF: Pair-feeding
PKA: Protein Kinase A

